# Loss-of-function variants in the cardiac K_v_11.1 channel as a genetic biomarker for SUDEP

**DOI:** 10.1101/2021.03.19.436102

**Authors:** Ming S. Soh, Richard D. Bagnall, Mark F. Bennett, Lauren E. Bleakley, Erlina S. Mohamed Syazwan, A. Marie Phillips, Mathew D.F. Chiam, Chaseley E. McKenzie, Michael Hildebrand, Douglas Crompton, Melanie Bahlo, Christopher Semsarian, Ingrid E. Scheffer, Samuel F. Berkovic, Christopher A. Reid

**Affiliations:** The Florey Institute of Neuroscience and Mental Health, University of Melbourne, Parkville, VIC, Australia; Agnes Ginges Centre for Molecular Cardiology at Centenary Institute, The University of Sydney, Sydney, NSW, Australia; Faculty of Medicine and Health, The University of Sydney, NSW, Australia; Population Health and Immunity Division, The Walter and Eliza Hall Institute of Medical Research, Melbourne, VIC, Australia; Department of Medical Biology, University of Melbourne, Melbourne, VIC, Australia; Epilepsy Research Centre, Department of Medicine, University of Melbourne, Austin Health, Heidelberg, VIC, Australia; School of Biosciences, University of Melbourne, Melbourne, VIC, Australia; Murdoch Children’s Research Institute, The Royal Children’s Hospital, Parkville, VIC, Australia; Neurology Department, Northern Health, Epping, VIC, Australia; Department of Paediatrics, University of Melbourne, Royal Children’s Hospital, VIC, Australia

**Keywords:** SUDEP, biomarkers, epilepsy, cardiac arrhythmia, genetics, ion channels, *KCNH2*, Kv11.1, common variants

## Abstract

**Objective:** To compare the frequency and impact on channel function of *KCNH2* variants in SUDEP patients with epilepsy controls comprising patients older than 50 years, a group with low SUDEP risk, and establish loss-of-function *KCNH2* variants as predictive biomarkers of SUDEP risk.

**Methods:** We searched for *KCNH2* variants with a minor allele frequency of < 5%. Functional analysis in *Xenopus laevis* oocytes was performed for all *KCNH2* variants identified.

**Results:** *KCNH2* variants were found in 11.1% (10/90) of SUDEP individuals compared to 6.0% (20/332) of epilepsy controls (*p* = 0.11). Loss-of-function *KCNH2* variants, defined as causing > 20% reduction in maximal amplitude, were observed in 8.9% (8/90) SUDEP patients compared to 3.3% (11/332) epilepsy controls suggesting about three-fold enrichment (nominal *p* = 0.04). *KCNH2* variants that did not change channel function occurred at a similar frequency in SUDEP (2.2%; 2/90) and epilepsy control (2.7%; 9/332) cohorts (*p* > 0.99). Rare *KCNH2* variants (< 1% allele frequency) associated with greater loss of function and an ∼11-fold enrichment in the SUDEP cohort (nominal *p* = 0.03). *In silico* tools were unable to predict the impact of a variant on function highlighting the need for electrophysiological analysis.

**Conclusions:** These data show that loss-of-function *KCNH2* variants are enriched in SUDEP patients and suggest that cardiac mechanisms contribute to SUDEP risk. We propose that genetic screening in combination with functional analysis can identify loss-of-function *KCNH2* variants that could act as biomarkers of an individual’s SUDEP risk.

## Introduction

People with epilepsy have a two-to three-fold increased risk of premature mortality, with Sudden Unexpected Death in Epilepsy (SUDEP) the most common cause of death^1^. SUDEP occurs without warning, most frequently in young adults. Frequent tonic-clonic seizures are the biggest risk factor for SUDEP^2-4^. Other risk factors are markers of seizure severity including epilepsy duration and poly-therapy^3-5^. However, SUDEP also occurs in patients with mild, well-controlled epilepsy, suggesting other risk factors exist^6^.

The pathophysiological mechanism(s) responsible for SUDEP are unclear. A systematic retrospective analysis of ten SUDEP deaths in the Incidence and Mechanisms of Cardiorespiratory Arrests in Epilepsy Monitoring Units (MORTEMUS) revealed that seizure-mediated terminal apnea always preceded terminal asystole^7^. The patient cohort in this study was small and involved individuals who were undergoing long-term video-EEG monitoring, implying refractory epilepsy. These cases may therefore not be representative of all individuals with SUDEP. Aside from seizure severity, the presence of abnormal cardiac rhythms has been widely implicated in SUDEP risk^8, 9^. Both human and animal studies show that seizure-mediated changes in cardiac electrophysiology occur, including seizure-driven cortical autonomic dysfunction and longer-term altered cardiac ion channel expression^10^. Furthermore, genetic studies have found variants in genes associated with cardiac arrhythmia syndromes in SUDEP cases^9, 11-14^, including genes that cause long QT syndrome (LQTS)^12-14^.

LQTS results from delayed myocardial repolarisation that manifests as a prolonged QT interval on the electrocardiography increasing the risk of ‘torsades de pointes’ that can trigger sudden cardiac death^15^. Here we focus on *KCNH2*, which was among the top 30 genes identified in a gene-based rare variant collapsing analysis of SUDEP patients and matched ancestry controls^12^. *KCNH2* encodes the pore-forming α subunit of the voltage-gated potassium channel K_v_11.1. *In vitro* screening assays have established that loss of K_v_11.1 function leads to familial LQTS type 2 (LQTS2)^16-18^. These same assays routinely assess modulation of K_v_11.1 by drug candidates, with block a strong predictor of cardiac toxicity^19^. *KCNH2* variants have been identified in SUDEP patients, including both rare pathogenic and common variants although statistically significant enrichment has not been demonstrated^12, 13, 20^. We have proposed that when combined, seizures and a risk variant in an arrhythmogenic gene could interact to significantly increase SUDEP risk^21^. This hypothesis predicts that SUDEP individuals will have an enrichment of *KCNH2* variants that impact channel function compared to epilepsy patients at low risk of SUDEP. To test this hypothesis, we measured the K_v_11.1-generated current of *KCNH2* variants found in SUDEP patients. The results were compared to *KCNH2* variants identified in a control sample comprising epilepsy patients who were over 50 years of age. Death rate is reported to be 6 times higher between the ages of 16 and 24 in epilepsy patients, with SUDEP accounting for ∼40% of deaths under the age of 45^22^. Our epilepsy control cohort is therefore considered to have ‘escaped’ SUDEP^23^. An enrichment of loss-of-function *KCNH2* variants in SUDEP compared to epilepsy controls argue that cardiac mechanisms may contribute to risk.

## Methods

### Identification of SUDEP and control variants

Our combined cohorts of 90 unrelated SUDEP patients were recruited from the epilepsy genetics research program in Melbourne, Australia, during life, or from coronial cases investigated at the Departments of Forensic Medicine in New South Wales, Victoria, Queensland, and South Australia, as previously described^24, 25^. All exons of *KCNH2* were Sanger-sequenced in 29 SUDEP cases, and the remaining 61 SUDEP cases underwent exome sequencing as previously described^12^. We looked for variants in the KCNH2 protein coding regions and essential splice site dinucleotides with an allele frequency < 5% in the gnomAD reference population database^26^. Variants identified by exome sequencing were Sanger-verified.

*KCNH2* variants were identified in our ‘epilepsy control’ cohort of 332 Australian individuals with generalised or non-lesional focal epilepsy over the age of 50 years (born before 1970) who underwent whole exome sequencing as part of our contribution to the Epi25 Consortium^27^. This population is drawn from a broad spectrum of epilepsy patients. Given that SUDEP is far more likely to occur between the ages of 20-40 years of age^28^, with death rate reported to be six times higher between the ages of 16 and 24 in epilepsy patients and accounting for ∼40% of deaths under the age of 45^29^, these patients were considered to have ‘escaped’ SUDEP, providing a suitable comparison population. Variant calling was performed as previously described^27^. Variants in the 332 individuals with epilepsy over the age of 50 were annotated using ANNOVAR^27^ and then filtered to obtain a list of single-nucleotide variants in *KCNH2* with an allele frequency < 5% in gnomAD^26^.

Ancestry predictions for the exome-sequenced SUDEP and epilepsy controls inferred 95% of samples in each cohort to be European. Exome sequencing achieved mean coverage greater than 20x across 78% of the *KCNH2* coding region in the SUDEP cohort and 94% in the epilepsy controls.

### Standard protocol approvals, registrations, and patient consents

The study was approved by the Human Research Ethics Committees of Austin Health and Royal Prince Alfred Hospital. Signed consent was provided by patients themselves, their parent, next-of-kin, or legal guardian in the case of children or patients with intellectual disability. Some of SUDEP cases were analysed in a de-identified manner. Some samples from control cohort had been collected over a 20-year period in some centers, so the consent forms reflected standards at the time of collection. Samples were only accepted if the consent did not exclude data sharing.

### *KCNH2* site-directed mutagenesis and *in vitro* cRNA preparation

cDNA encoding a full-length transcript (NM_000238.4 Ensembl database) of human *KCNH2* was subcloned into the *Xenopus* oocyte expression vector pGEMHE-MCS. Site-directed mutagenesis to create human *KCNH2* variants was completed by Genscript Biotech (Piscataway, NJ, USA). All clones were verified by Sanger sequencing. *In vitro* synthesis of cRNA was performed using linearised cDNA template and the mMessage mMachine® T7 transcription kit (Ambion, Thermo Fisher Scientific, Waltham, MA). RNA integrity was assessed both spectrophotometrically (Nanodrop) and by gel electrophoresis. All cRNAs were stored at −80°C.

### Oocyte extraction

Adult female *Xenopus laevis* frogs were housed at the Florey Institute of Neuroscience and Mental Health. Animal procedures and oocyte preparation followed standard procedures in accordance with the conditions approved by the Florey’s Ethics Committee. Briefly, frogs were anesthetised with 1.3 mg/ml tricaine methanesulfonate and oocytes were surgically removed via a small incision to the abdomen. Oocytes were defolliculated with 1.5 mg/ml collagenase for two hours and rinsed with OR-2 solution (in mM: 82.5 NaCl, 2 KCl, 1 MgCl_2_.6H_2_O, 5 HEPES, pH 7.4). Healthy mature oocytes stage V or VI were isolated.

### Channel expression

The NM_000238.4 *KCNH2* transcript was set as our control sequence and designated as the wild-type (WT) channel. 0.5 ng of cRNAs (in 50 nl) coding for *KCNH2* were manually injected into the oocytes. Injected oocytes were maintained in ND96 storage solution (in mM: 96 NaCl, 2 KCl, 1 MgCl_2_.6H_2_O, 1.8 CaCl_2_.2H_2_O, 5 HEPES, 50 mg/l gentamicin, pH 7.4) at 17 °C for two days to allow translation and trafficking of channels prior to recording.

### Two-electrode voltage clamp electrophysiology

Standard two-electrode voltage clamp hardware was used (TEC-05X or TEC10X, NPI, Tamm, Germany). Oocytes were impaled with microelectrodes with an input resistance of between 0.2-2.0 MΩ, containing 3 M KCl. During experiments oocytes were continually perfused with high K^+^ solution (in mM: 100 KCl, 1.8 CaCl_2_, 1 MgCl_2_, 10 HEPES, pH 7.4) and clamped at −90 mV. Incremental 10 mV voltage steps from −90 to +50 mV were applied for 0.5 s, followed by a test pulse at −100 mV for 3.2 s. Data were sampled at 5000 Hz and low-pass-filtered at 1 kHz. Voltage clamp control and data acquisition were obtained using pCLAMP v8.10 software (Molecular Devices, USA). All recordings were performed at 18-20°C. Currents from WT channels were always measured contemporaneously using the same batch of oocytes.

### Electrophysiology analysis

Raw currents were baseline-adjusted and leak-subtracted, and peak current amplitudes were analysed offline using AxoGraph v1.7.4 (AxoGraph Scientific, Sydney, AU). Half-maximal activation voltage and slope values were obtained from current-voltage curves that were fitted with the Boltzmann equation after normalising the peak test pulse current after each voltage step to the maximum peak test pulse current. Wild-type-normalised current-voltage curves were obtained by normalising peak test pulse current after each voltage step to the average maximum wild-type peak test pulse current and fitted with the Boltzmann equation.

### Statistical analysis

#### Functional analysis

An F-test was used to check for variance equality. Standard one-way ANOVA with Dunnett’s post-hoc correction was used for statistical comparison to wild-type values if variances were approximately equal (*p* > 0.05). Welch’s ANOVA with Dunnett’s post-hoc correction was used instead, for unequal variances (*p* < 0.05). Significance was set at uncorrected *p* < 0.05 for ANOVA tests. Statistical analysis for association used the Fisher’s Exact Test. Graphs and statistical tests were generated and performed using Prism v8.1.0 (GraphPad, CA, USA). All data points are shown as mean ± S.E.M.

### Data availability

Data collected for the study will be made available upon reasonable request to the corresponding author.

## Results

### *KCNH2* variants in SUDEP patients and a control epilepsy population

*KCNH2* variants previously described in our SUDEP cohorts are listed in Table 1. Ten out of 90 individuals suffering SUDEP carried a missense or truncation *KCNH2* variant with < 5% minor allele frequency^12, 25^. These variants have a range of pathogenicity classifications according to the ClinVar database^30^. R744X is reported as pathogenic; Y54H, G924A, R176W, and R1047L are variants of uncertain clinical significance; while G749A is not reported in the ClinVar database.

**Table 1:**
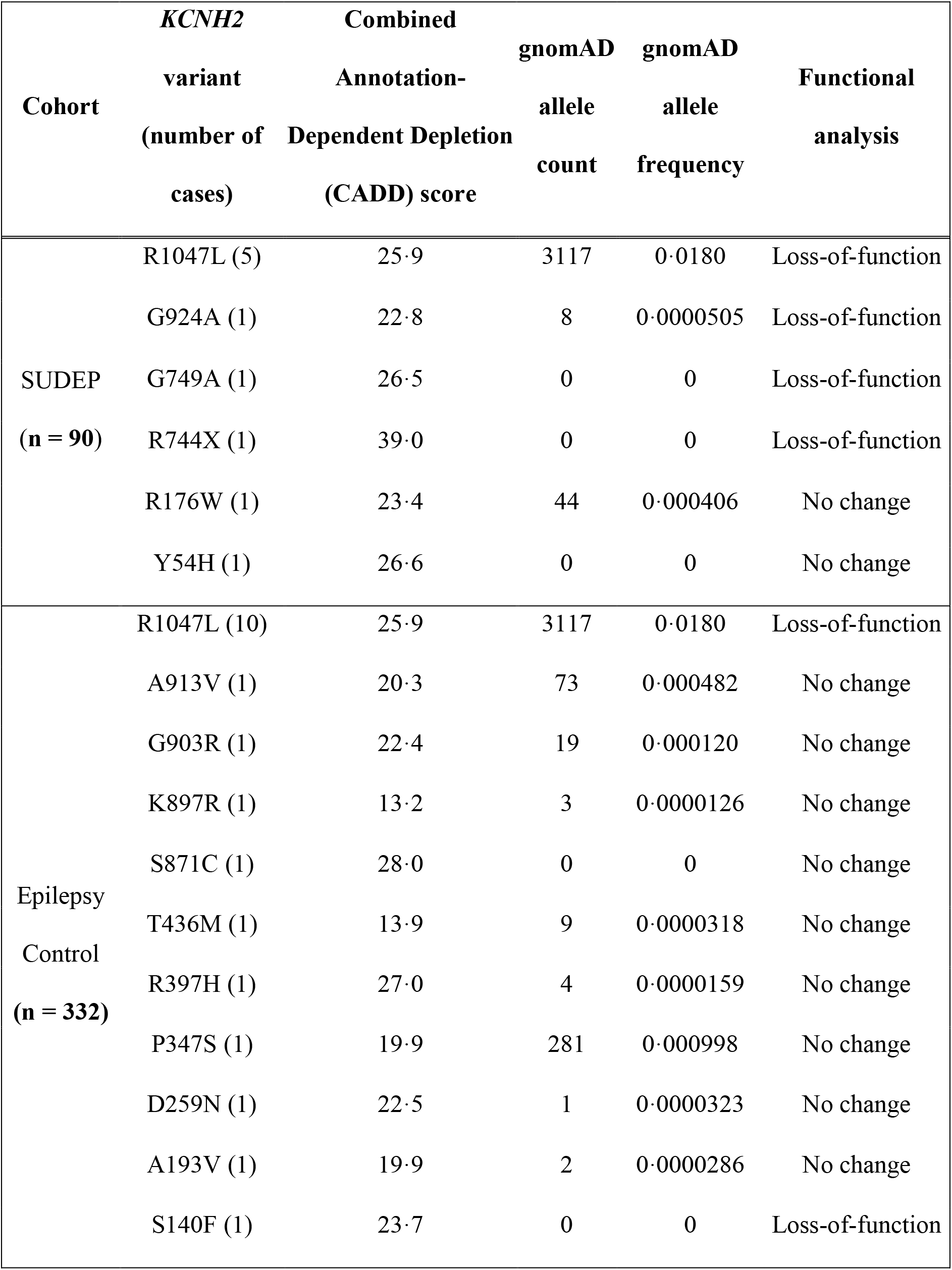
*KCNH2* variants found in SUDEP and epilepsy control populations

In our epilepsy control population of 332 living patients with epilepsy over 50 years of age with whole exome sequencing data, variants in *KCNH2* that satisfied the filtering criteria were found in 20 out of 332 subjects (Table 1). These included the common R1047L variant also found in SUDEP cases (Table 1). Variants A913V, K897R, S871C, T436M, R397H, P347S, and D259N were classified as variants of uncertain significance in ClinVar, while A193V and S140F are not reported in ClinVar.

*KCNH2* variants were found in 11.1% of SUDEP cases (10/90) compared to 6.0% of epilepsy controls (20/332; *p* = 0.11).

### *In silico* predictions of K_v_11.1 channel dysfunction and SUDEP risk

*In silico* prediction tools provide a method of estimating the detrimental impact of a given variant on protein function. We determined the Combined Annotation Dependent Depletion (CADD) score for each *KCNH2* variant (Table 1). Only the *KCNH2* R744X variant had a CADD score above 30 indicating, as in ClinVar, that it was pathogenic.

### Functional characterisation of *KCNH2* variants

Manual two-electrode voltage clamp was used to record currents from wild-type (WT; NM_000238.4 *KCNH2* transcript) and mutated K_V_11.1 channels expressed in *Xenopus* oocytes (Figure 1). Channels were activated by a series of depolarising voltage excursions from a holding potential of −90 mV. Maximal tail currents were used to measure channel activity. In our SUDEP cohort, functional analysis revealed a significant reduction in current amplitude (> 20%) for R1047L, G924A, G749A, and R744X mutated channels relative to wild-type channels (Figures 1A-E, H). R176W, and Y54H variants had no effect (Figures 1F-H). In the epilepsy control group, the R1047L (also in the SUDEP cohort) and the S140F variant reduced current amplitude (Figure 2). The other nine epilepsy control variants were without effect on K_V_11.1 channel current amplitude (Figure 2). Other biophysical parameters measured, including the half maximal activation voltage and Boltzmann slope, are reported in Figure 3. In the SUDEP group, only G924A and G749A showed altered biophysical properties (Figures 3A, C), whereas R744X could not be measured. No variants in the epilepsy control group showed any differences in other biophysical parameters (Figures 3B, D). Changes in biophysical properties for SUDEP *KCNH2* variants may contribute to overall channel dysfunction and consequently increase risk of sudden death.

**Figure 1:**
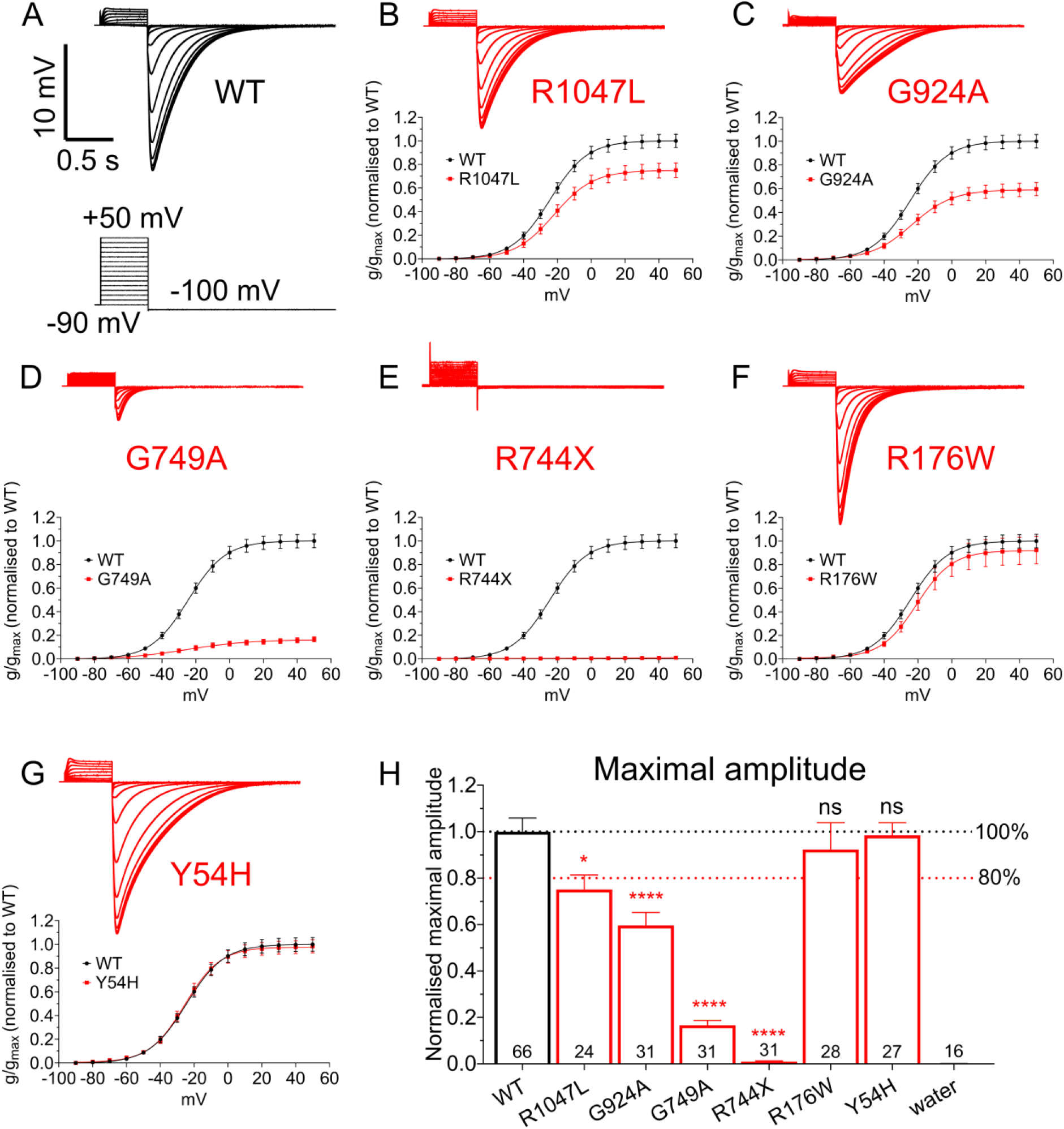
Functional analysis of *KCNH2* variants from SUDEP patients. **(A)** Sample recording traces of K_v_11·1 wild-type (WT) channels. *Insert:* cartoon of the voltage protocol applied. **(B)** Sample recording traces of K_v_11·1 variant channels (*top*) and average normalised conductance-voltage relationships (*below*) comparing K_v_11·1 WT and variant channels for **(B)** R1047L, **(C)** G924A, **(D)** G749A, **(E)** R744X, **(F)** R176W, and **(G)** Y54H variants. **(H)** Average maximal amplitude for each variant. Number in each bar represent the number of independent oocytes recorded for each variant. Black and red dashed lines indicate 100% and 80% respectively of maximal current amplitude of K_v_11·1 WT channel. * *p* < 0·05, **** *p* < 0·0001.

**Figure 2:**
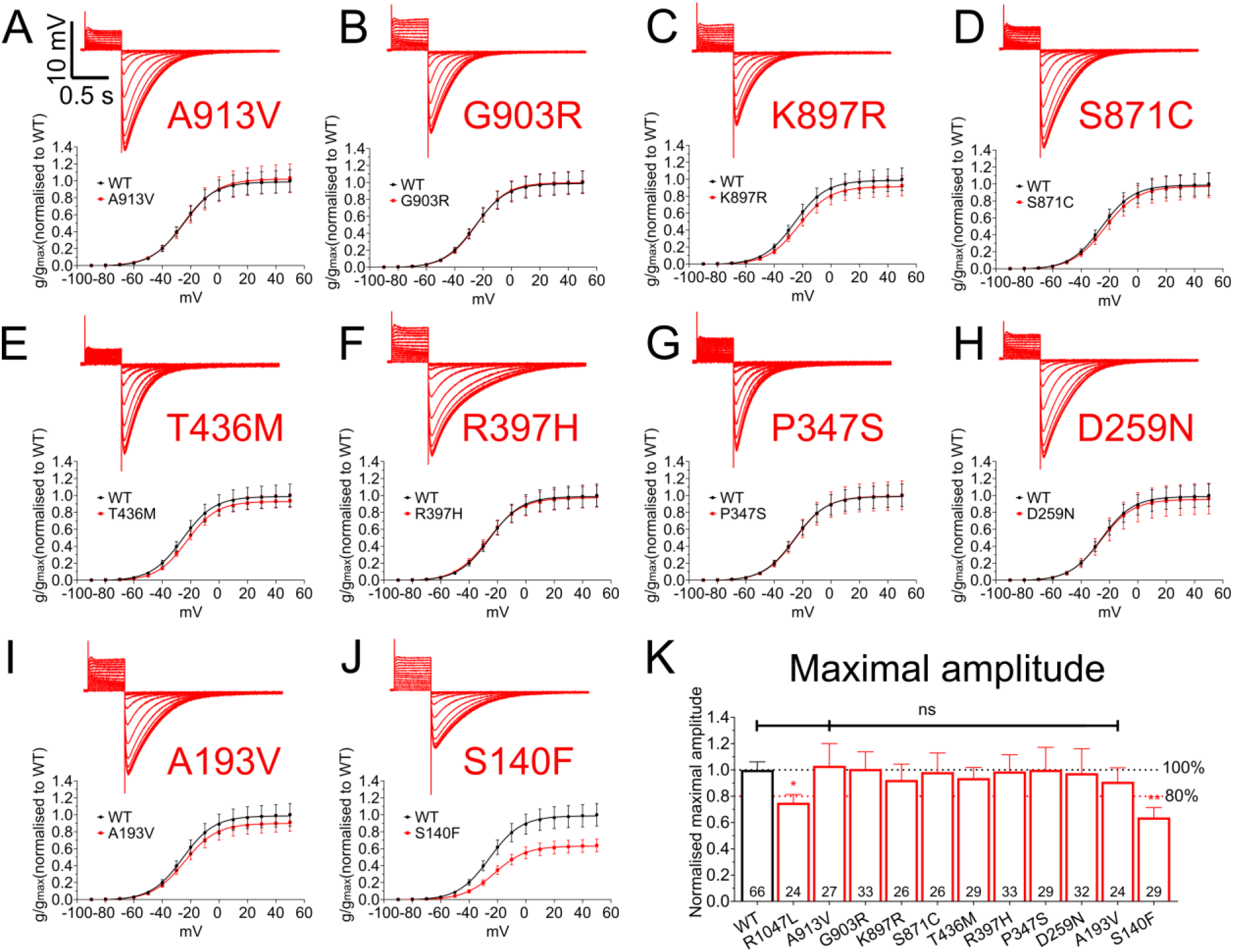
Functional analysis of *KCNH2* variants from epilepsy control cohort. Sample recording traces of K_v_11·1 variant channels (*top*) and average normalised conductance-voltage relationships (*below*) comparing K_v_11·1 WT and variant channels for **(A)** A913V **(B)** G903R, **(C)** K897R, **(D)** S871C, **(E)** T436M, **(F)** R397H, **(G)** P347S, **(H)** D259N **(I)**, A193V, and **(J)** S140F variants. **(K)** Average maximal amplitude for each variant in epilepsy control cohort. Number in each bar represent the number of independent oocytes recorded for each variant. Black and red dashed lines indicate 100% and 80% respectively of maximal current amplitude of the K_v_11·1 WT channel. * *p* < 0·05, ** *p* < 0·01.

**Figure 3:**
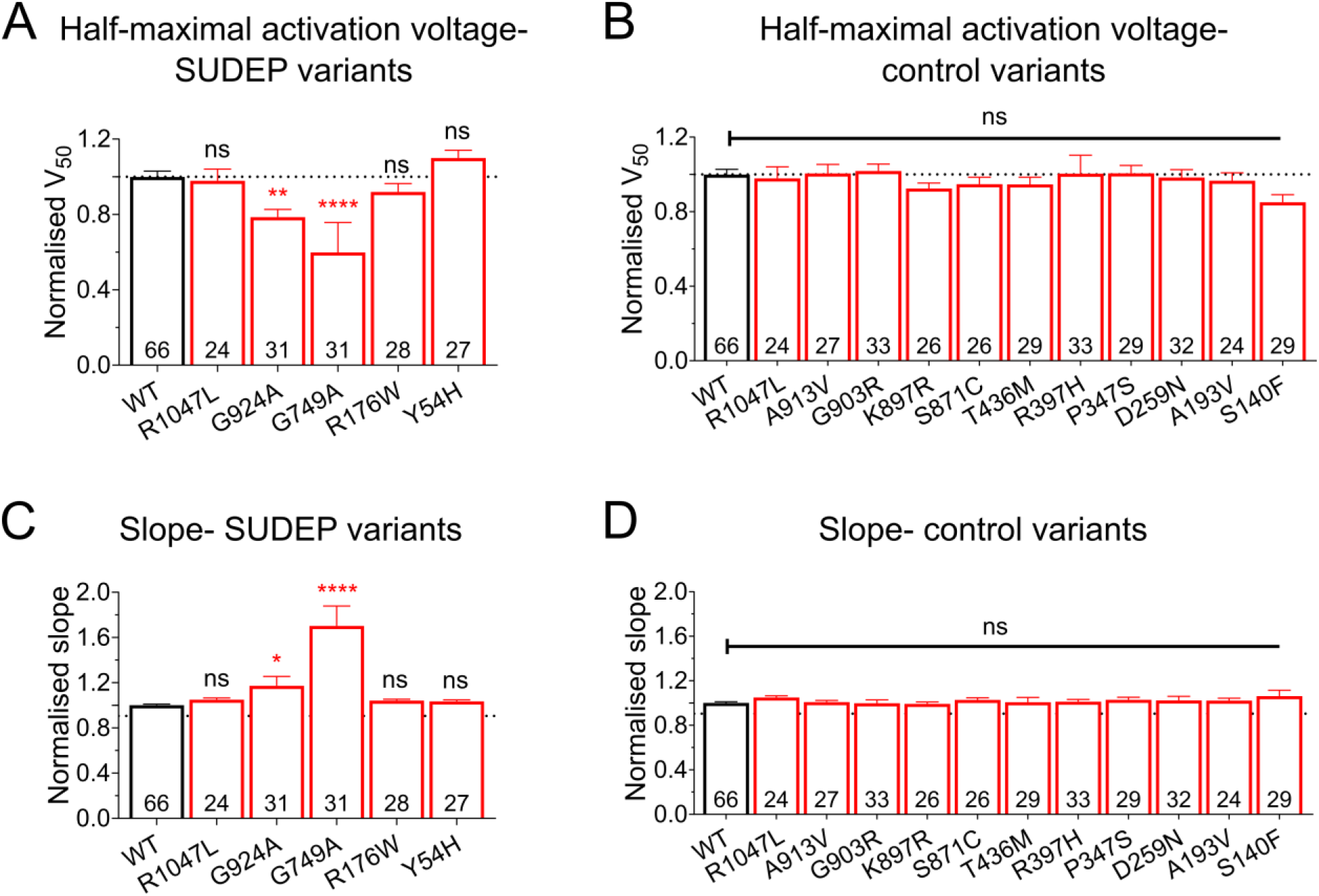
Biophysical properties of *KCNH2* variants from SUDEP cases and epilepsy control population. **(A)** Average half-maximal voltage of activation for each variant in SUDEP cohort. **(B)** Average half-maximal voltage of activation for each variant in epilepsy control cohort. **(C)** Average slope from the Boltzmann fit for each variant in the SUDEP cohort. **(D)** Average slope from the Boltzmann fit for each variant in the epilepsy control cohort. Number in each bar represent the number of independent oocytes recorded for each variant. * *p* < 0·05, ** *p* < 0·01, **** *p* < 0·0001.

### Enrichment of loss-of-function *KCNH2* variants in SUDEP cases

Our functional data allows classification of each variant as either loss-of-function, defined as a statistically significant reduction in current amplitude of > 20%, or no change in function (Table 1). Based on these criteria, eight out of 90 (8.9%) SUDEP cases carried a *KCNH2* loss-of-function variant, while two out of 90 (2.2%) carried a variant in which function was not changed. In contrast, our epilepsy patient control population had 11 out of 332 (3.3%) patients with a loss-of-function variant, and nine out of 332 (2.7%) carried a variant which did not alter function. The SUDEP cohort has approximately three-fold enrichment for loss-of-function *KCNH2* variants (OR = 2.7, 95% confidence interval (1.1, 7.4), Fisher’s exact test nominal *p* = 0.04) compared with the epilepsy control population (Figure 4A). There was no enrichment of *KCNH2* variants that did not change channel function in the SUDEP cohort compared to the epilepsy control cohort (OR = 0.8, 95% confidence interval 0.17 to 3.8, Fisher’s exact *p* > 0.99).

**Figure 4:**
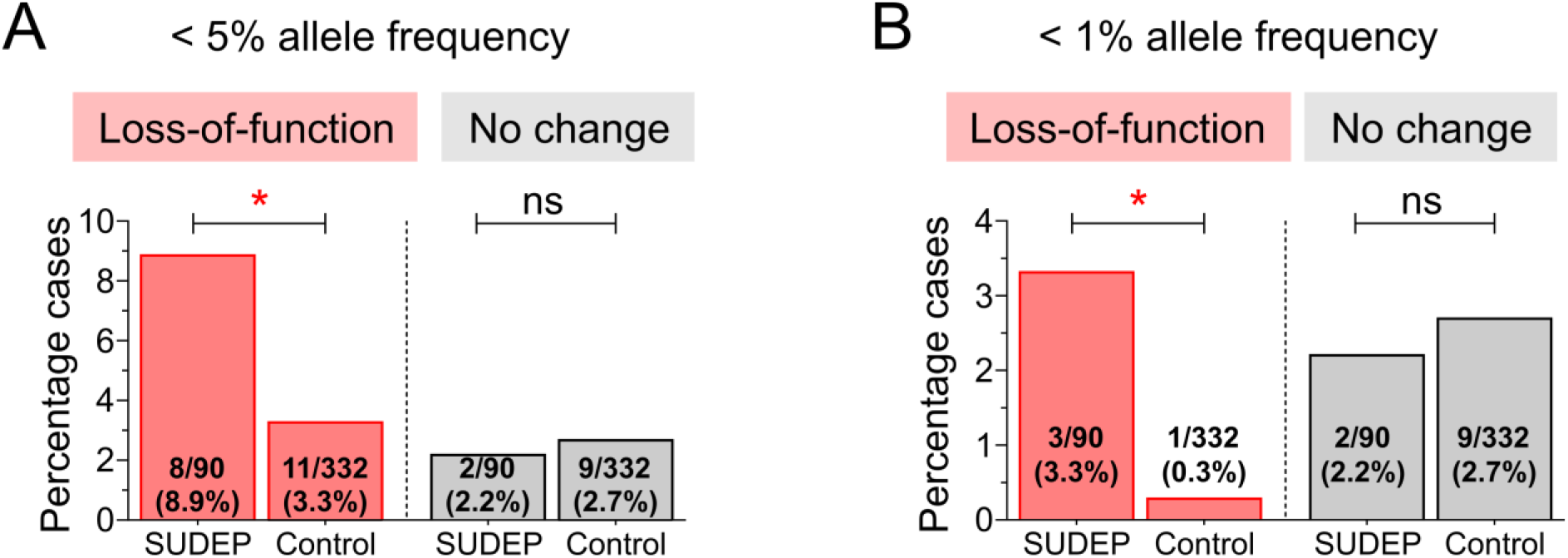
Enrichment of *KCNH2* variants in SUDEP and epilepsy control cohorts. **(A)** *KCNH2* variants with less than 5% allele frequency are enriched approximately three times in SUDEP compared to epilepsy control cohort. **(B)** Enrichment of rare *KCNH2* variants with less than 1% allele frequency is further increased to about ten times in SUDEP cohort. * *p* < 0·05.

In the SUDEP cohort, rare *KCNH2* variants with a minor allele frequency of < 1% associate with greater loss of function implicating increased risk (Figure 1). Consistent with this, rare *KCNH2* variants are enriched with a greater odds ratio (OR = 11.4, 95% confidence interval 1.2 to 111.1, Fisher’s exact *p* = 0.03) (Figure 4B). R1047L is the most common variant that we tested. It is observed in approximately 3.5% of the general population^26^ and was found in five out of 90 (5.6%) SUDEP cases and ten out of 332 (3.0%) epilepsy control patients. The reduction in current amplitude for R1047L variant shows that loss-of-function is not confined to rare *KCNH2* variants.

## Discussion

Previous genetic screening efforts in SUDEP have identified rare variants in genes that are associated with severe developmental and epileptic encephalopathies^31^. This is to be expected, as these variants cause severe epilepsies which carry a high SUDEP risk^31, 32^. Whether other genetic risk factors contribute to SUDEP risk is less clear. Screening efforts in SUDEP cases have identified variants in cardiac genes that cause arrhythmia syndromes^12, 25, 33-36^. We have proposed that when combined, seizures and a risk variant in an arrhythmogenic gene could interact to significantly increase SUDEP risk^37^. Here we focused on *KCNH2*, in which loss-of-function variants are an established cause of LQTS leading to sudden death. We show an approximate three-fold enrichment in loss-of-function *KCNH2* variants in our SUDEP cohort relative to an older epilepsy control cohort that has ‘escaped’ SUDEP. These data support the premise that *KCNH2* loss-of-function variants act as genetic biomarkers of SUDEP risk and motivates the need to examine this hypothesis in additional, independent SUDEP cohorts.

The pathophysiological mechanism(s) responsible for SUDEP are likely to be multifactorial. A systematic retrospective analysis of ten SUDEP deaths in the Incidence and Mechanisms of Cardiorespiratory Arrests in Epilepsy Monitoring Units (MORTEMUS) revealed that seizure-mediated terminal apnea always preceded terminal asystole^38^. This is strong evidence implying respiratory factors as a cause of death. However, the patient cohort in this study was small and involved individuals who were undergoing long-term video-EEG monitoring, implying refractory epilepsy. These cases may therefore not be representative of all individuals with SUDEP. Our data suggests that cardiac factors are important, at least in a subset of SUDEP cases.

Characterisation of *KCNH2* variants based on *in silico* predictions of protein dysfunction using the CADD method were uninformative. Only the truncation variant, R744X, had a CADD score greater than 30 and is therefore predicted to be deleterious. CADD scores for the G924A and G749A variants were relatively low yet functional analysis in *Xenopus* oocytes revealed significant reductions in current amplitude. As noted below, functional analysis methods are not without limitation. However, our findings highlight that functional analysis should remain the gold standard by which to judge potential pathogenicity until *in silico* prediction tools improve.

In this study we have categorised *KCNH2* variants into either loss-of-function or no change in function. However, individual variants do vary in the degree of functional impairment shown *in vitro* and thus are unlikely to contribute equally to the risk of sudden death. In the case of rare pathogenic variants, such as R744X, G749A and G924A there is a large loss-of-function observed in the expression assay and we infer a stronger likelihood that cardiac arrhythmia contributed to death. Increased enrichment of rare *KCNH2* variants (OR=11.4) in the SUDEP cohort is consistent with a correlation between the extent of loss of function and increased risk. More common variants with lesser *in vitro* functional impairment may contribute less individual risk. For example, R1047L has an allele frequency of 1.8% in the gnomAD database of population controls and has a smaller impact on channel function. The R1047L variant is likely to increase individual risk less when compared to variants that cause a large impact on channel function. However, at the population level the R1047L variant impact is likely to be significant given its common nature, increasing risk to a small degree in many people. Additional studies looking at the functional impact of *KCNH2* variants in a greater number of SUDEP patients will be required to fully understand attributable risk. It is also important to note that heterologous expression systems cannot report on more complex cell function. More sophisticated model systems such as cardiac myocytes derived from stem cells will help further define the relationship between *KCNH2* variants and arrhythmia risk.

There is surprisingly little evidence that directly links acute seizures to genetically-caused cardiac arrhythmia and sudden death. The Kcnq1 T311I mouse model of LQTS provides some evidence with over half of the recorded abnormalities in cardiac rhythm associated with epileptiform discharges^39^. This has implications for more common variants, such as R1047L, that are unlikely to be pathogenic alone but may increase risk of death in the context of seizures. Furthermore, both human and animal studies show that seizure-mediated changes in cardiac electrophysiology occur^40^. This includes seizure-driven dysfunction that can alter acute cardiac rhythm^41, 42^. Studies have also observed altered cardiac ion channel expression with ongoing seizure activity^43^. Either acute and/or chronic changes in cardiac function due to seizures may increase susceptibility to cardiac arrhythmias in people with a ‘loss-of-function’ *KCNH2* variant to significantly increase SUDEP risk.

The ability to identify patients at risk of SUDEP has important clinical implications. In patients with epilepsy carrying loss-of-function *KCNH2* variants, prolonged electroencephalogram with cardiac monitoring might be informative to explore ictal and interictal changes in cardiac rhythm, while prolonged cardiac monitoring with cardiac loop recorders could be considered for more prolonged interrogation of cardiac rhythm. This would allow the detection of additional biomarkers of risk, including arrhythmogenic markers such as prolonged QT intervals, especially during seizures. Patients identified to be at risk of cardiac arrhythmia could start prophylactic treatment with beta blockers which are used effectively in LQTS. Drugs known to impact QT intervals should also be avoided in such epilepsy patients.

*KCNH2* variants will only ever be one biomarker that will be part of a risk assessment. Seizure severity and frequency remain significant predictors of SUDEP risk^31, 32^. Further investigation into other potential genetic biomarkers, including arrhythmogenic genes such as *KCNQ1* and *SCN5A* is required, as are studies into how acute seizures or long-term seizure-related changes in cardiac function interact with genetic causes of arrhythmia. An ultimate goal is to develop a SUDEP risk matrix integrating the various clinical, genetic and environmental factors, and to prevent SUDEP by targeting all modifiable risk factors.

In conclusion, our data provides evidence that both rare and more common *KCNH2* variants that cause loss-of-function may act as biomarkers of SUDEP in epilepsy patients. These data need to be replicated in larger independent study cohorts. Our data motivates more focused clinical studies investigating the impact of loss-of-function *KCNH2* variants on cardiac rhythm. Our study also motivates the development of more complex *in vitro* models, as well as animal models, that will allow the interaction between seizures and genetic cardiac abnormalities to be investigated. These models will also provide an opportunity to test novel therapeutic strategies for the prevention of SUDEP.

## Appendix 1: Authors

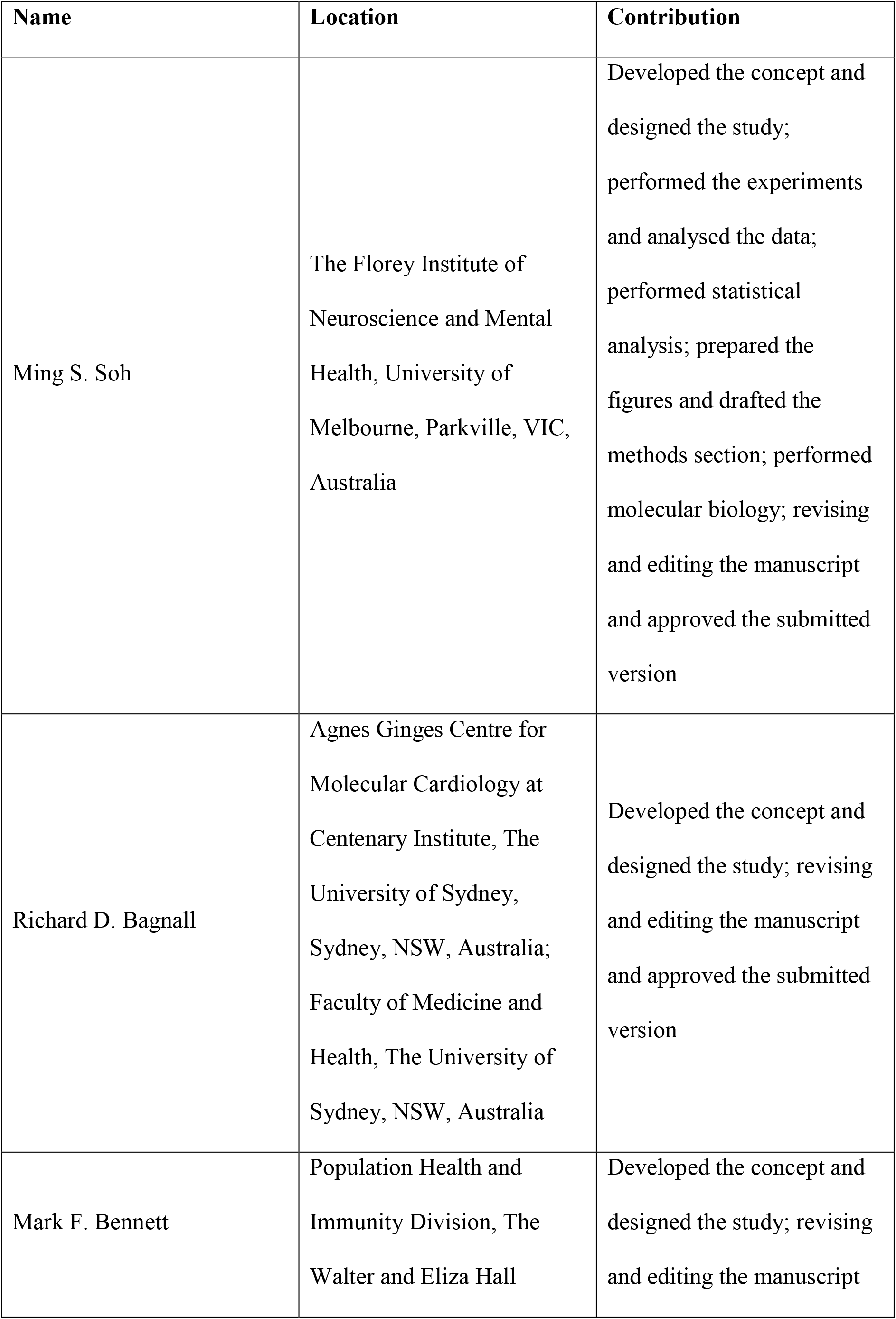
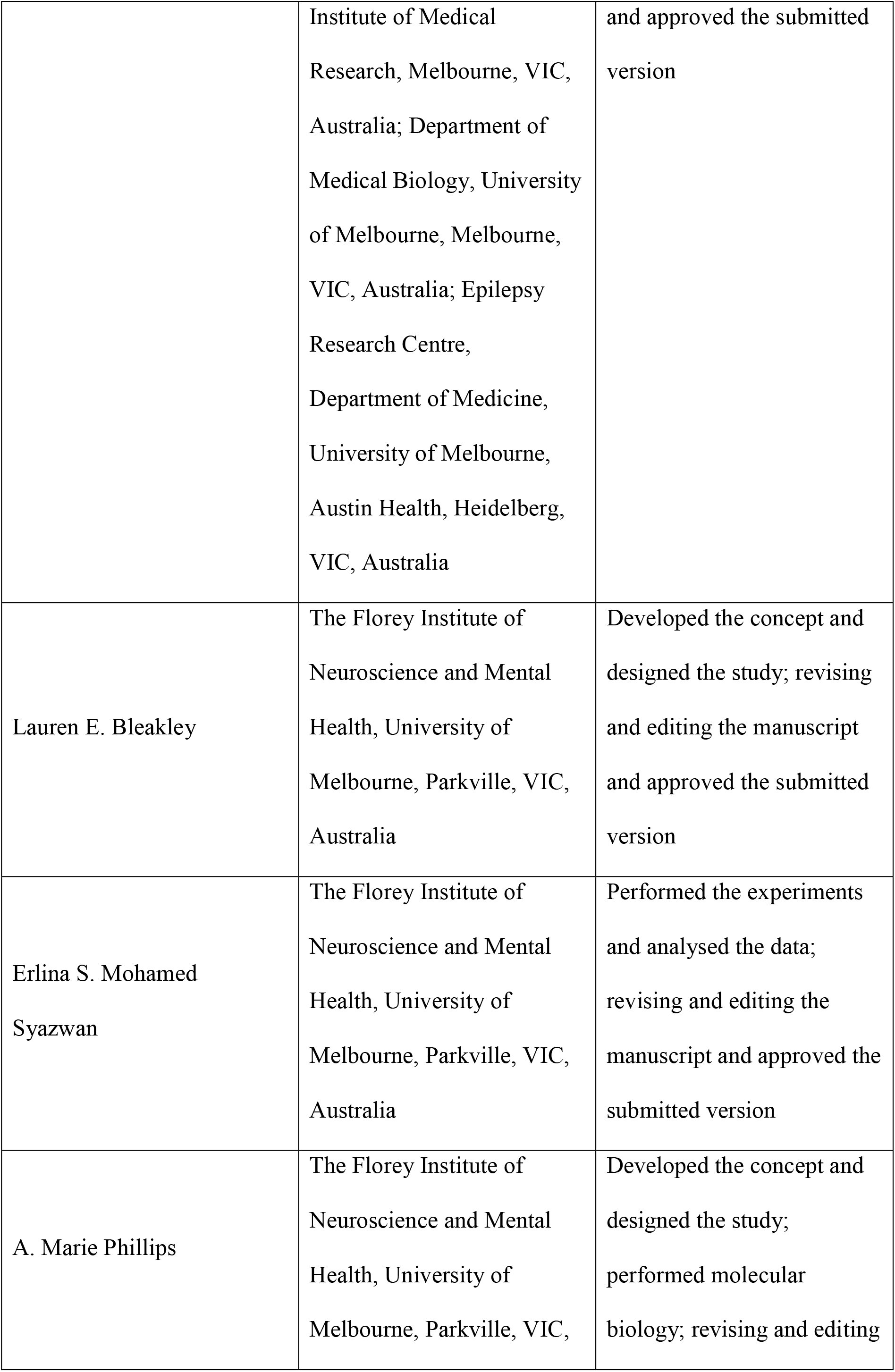
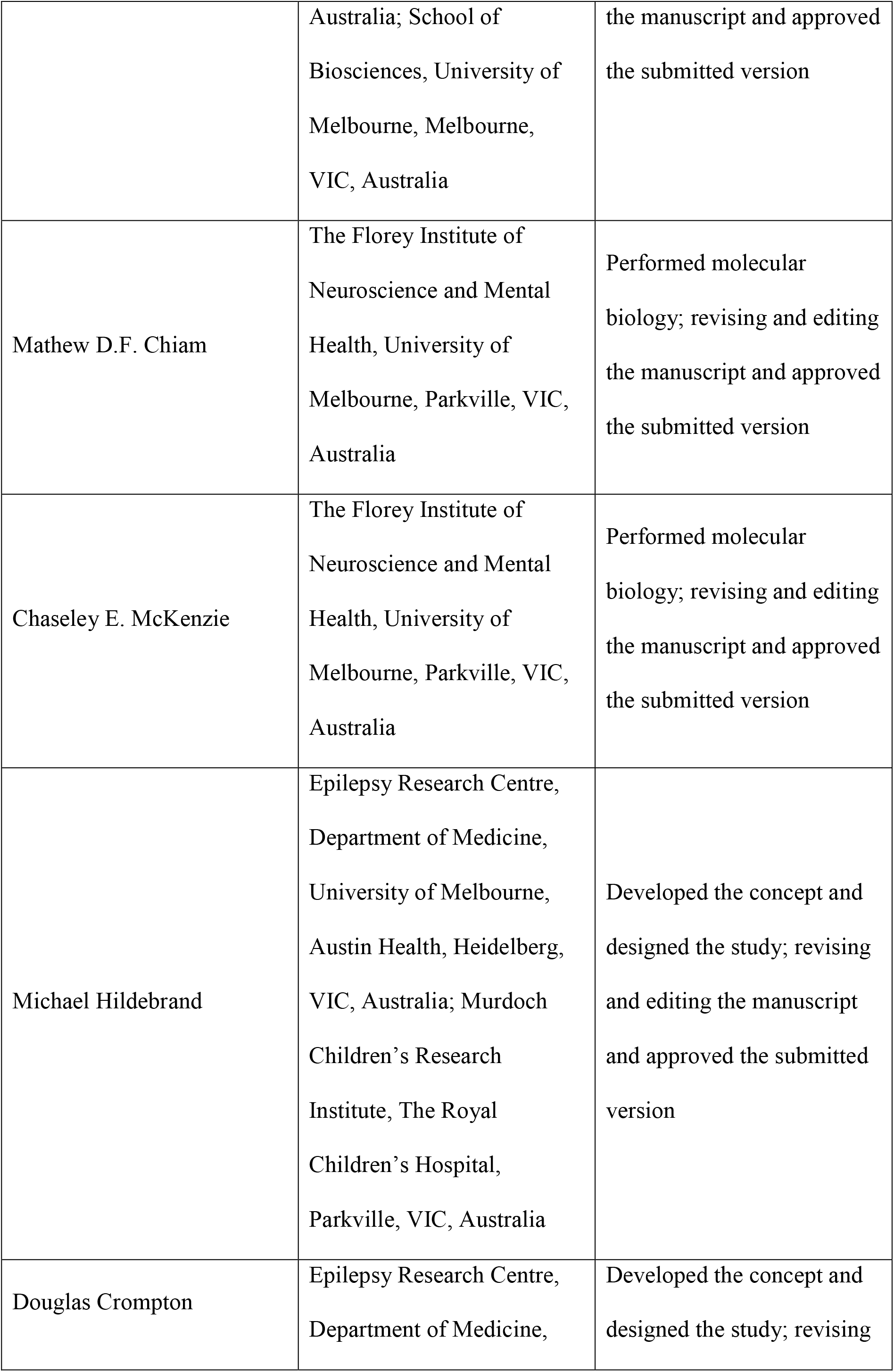
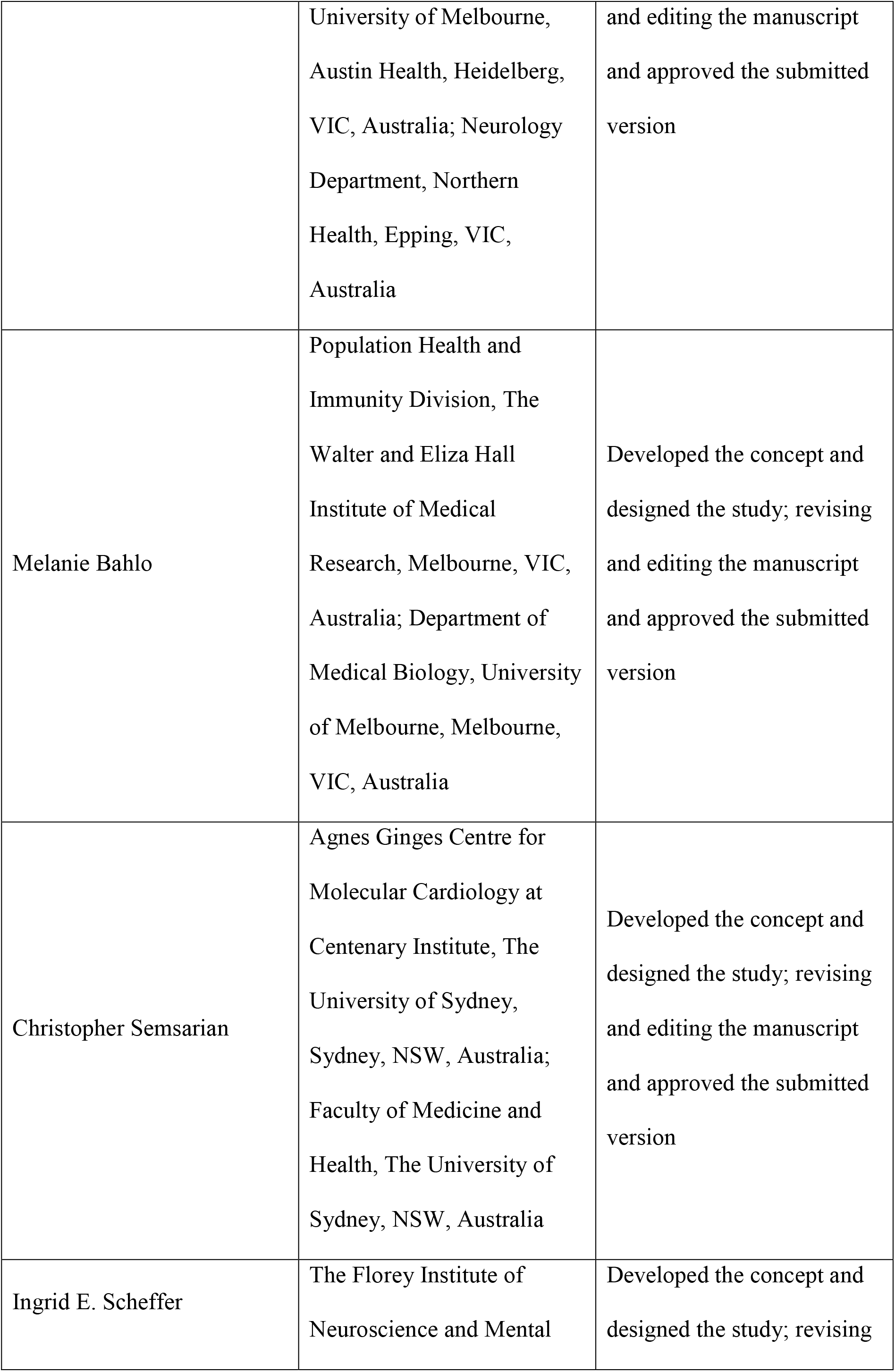
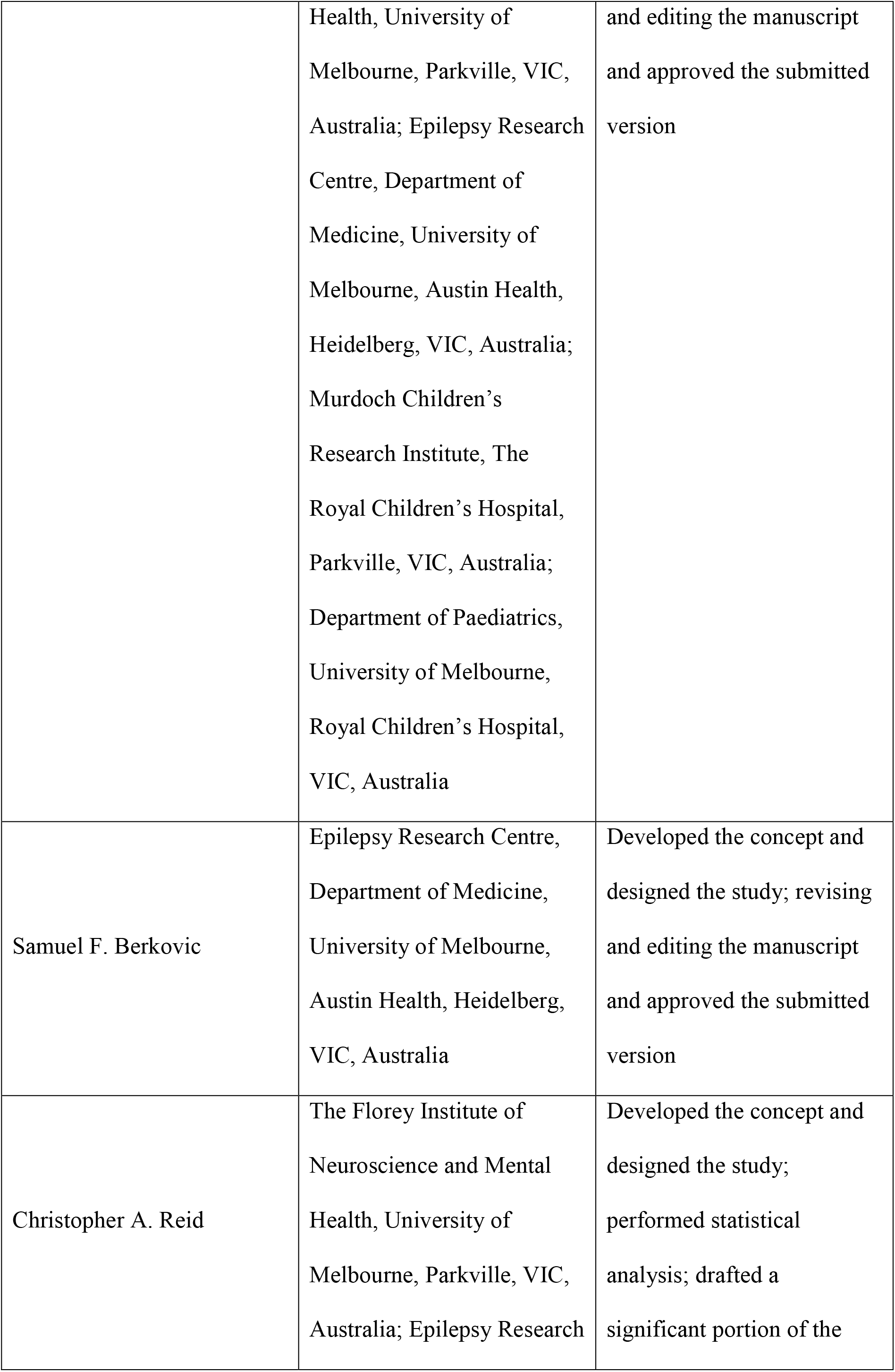
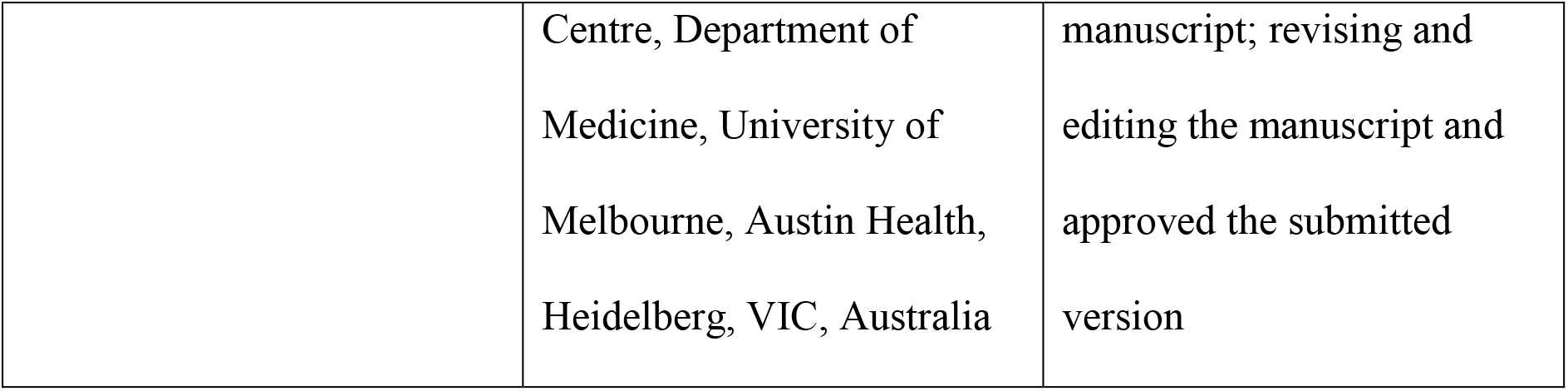

| **Name** | **Location** | **Contribution** |
| --- | --- | --- |
| Ming S. Soh | The Florey Institute of Neuroscience and Mental Health, University of Melbourne, Parkville, VIC, Australia | Developed the concept and designed the study; performed the experiments and analysed the data; performed statistical analysis; prepared the figures and drafted the methods section; performed molecular biology; revising and editing the manuscript and approved the submitted version |
| Richard D. Bagnall | Agnes Ginges Centre for Molecular Cardiology at Centenary Institute, The University of Sydney, Sydney, NSW, Australia; Faculty of Medicine and Health, The University of Sydney, NSW, Australia | Developed the concept and designed the study; revising and editing the manuscript and approved the submitted version |
| Mark F. Bennett | Population Health and Immunity Division, The Walter and Eliza Hall | Developed the concept and designed the study; revising and editing the manuscript |
|  | Institute of Medical Research, Melbourne, VIC, Australia; Department of Medical Biology, University of Melbourne, Melbourne, VIC, Australia; Epilepsy Research Centre, Department of Medicine, University of Melbourne, Austin Health, Heidelberg, VIC, Australia | and approved the submitted version |
| Lauren E. Bleakley | The Florey Institute of Neuroscience and Mental Health, University of Melbourne, Parkville, VIC, Australia | Developed the concept and designed the study; revising and editing the manuscript and approved the submitted version |
| Erlina S. Mohamed Syazwan | The Florey Institute of Neuroscience and Mental Health, University of Melbourne, Parkville, VIC, Australia | Performed the experiments and analysed the data; revising and editing the manuscript and approved the submitted version |
| A. Marie Phillips | The Florey Institute of Neuroscience and Mental Health, University of Melbourne, Parkville, VIC, | Developed the concept and designed the study; performed molecular biology; revising and editing |
|  | Australia; School of Biosciences, University of Melbourne, Melbourne, VIC, Australia | the manuscript and approved the submitted version |
| Mathew D.F. Chiam | The Florey Institute of Neuroscience and Mental Health, University of Melbourne, Parkville, VIC, Australia | Performed molecular biology; revising and editing the manuscript and approved the submitted version |
| Chaseley E. McKenzie | The Florey Institute of Neuroscience and Mental Health, University of Melbourne, Parkville, VIC, Australia | Performed molecular biology; revising and editing the manuscript and approved the submitted version |
| Michael Hildebrand | Epilepsy Research Centre, Department of Medicine, University of Melbourne, Austin Health, Heidelberg, VIC, Australia; Murdoch Children's Research Institute, The Royal Children's Hospital, Parkville, VIC, Australia | Developed the concept and designed the study; revising and editing the manuscript and approved the submitted version |
| Douglas Crompton | Epilepsy Research Centre, Department of Medicine, | Developed the concept and designed the study; revising |
|  | University of Melbourne,<br>Austin Health, Heidelberg,<br>VIC, Australia; Neurology<br>Department, Northern<br>Health, Epping, VIC,<br>Australia | and editing the manuscript<br>and approved the submitted<br>version |
| Melanie Bahlo | Population Health and<br>Immunity Division, The<br>Walter and Eliza Hall<br>Institute of Medical<br>Research, Melbourne, VIC,<br>Australia; Department of<br>Medical Biology, University<br>of Melbourne, Melbourne,<br>VIC, Australia | Developed the concept and<br>designed the study; revising<br>and editing the manuscript<br>and approved the submitted<br>version |
| Christopher Semsarian | Agnes Ginges Centre for<br>Molecular Cardiology at<br>Centenary Institute, The<br>University of Sydney,<br>Sydney, NSW, Australia;<br>Faculty of Medicine and<br>Health, The University of<br>Sydney, NSW, Australia | Developed the concept and<br>designed the study; revising<br>and editing the manuscript<br>and approved the submitted<br>version |
| Ingrid E. Scheffer | The Florey Institute of<br>Neuroscience and Mental | Developed the concept and<br>designed the study; revising |
|  | Health, University of Melbourne, Parkville, VIC, Australia; Epilepsy Research Centre, Department of Medicine, University of Melbourne, Austin Health, Heidelberg, VIC, Australia; Murdoch Children's Research Institute, The Royal Children's Hospital, Parkville, VIC, Australia; Department of Paediatrics, University of Melbourne, Royal Children's Hospital, VIC, Australia | and editing the manuscript and approved the submitted version |
| Samuel F. Berkovic | Epilepsy Research Centre, Department of Medicine, University of Melbourne, Austin Health, Heidelberg, VIC, Australia | Developed the concept and designed the study; revising and editing the manuscript and approved the submitted version |
| Christopher A. Reid | The Florey Institute of Neuroscience and Mental Health, University of Melbourne, Parkville, VIC, Australia; Epilepsy Research | Developed the concept and designed the study; performed statistical analysis; drafted a significant portion of the |
|  | Centre, Department of<br>Medicine, University of<br>Melbourne, Austin Health,<br>Heidelberg, VIC, Australia | manuscript; revising and<br>editing the manuscript and<br>approved the submitted<br>version |

## Declaration of interests

SFB declares unrestricted educational grants from UCB Pharma, SciGen and Eisai and consultancy fees from Praxis Precision Medicines. IES has served on scientific advisory boards for UCB, Eisai, GlaxoSmithKline, BioMarin, Nutricia, Rogcon and Xenon Pharmaceuticals; has received speaker honoraria from GlaxoSmithKline, UCB, BioMarin, Biocodex and Eisai; has received funding for travel from UCB, Biocodex, GlaxoSmithKline, Biomarin and Eisai; has served as an investigator for Zogenix, Zynerba, Ultragenyx, GW Pharma, UCB, Eisai, Anavex Life Sciences, Ovid Therapeutics, Epigenyx, Encoded Therapeutics and Marinus; and has consulted for Zynerba Pharmaceuticals, Atheneum Partners, Ovid Therapeutics, Epilepsy Consortium and UCB. IES may accrue future revenue on pending patent WO61/010176 (filed: 2008): Therapeutic Compound; has a patent for *SCN1A* testing held by Bionomics Inc and licensed to various diagnostic companies; has a patent molecular diagnostic/theranostic target for benign familial infantile epilepsy (BFIE) [PRRT2] 2011904493 & 2012900190 and PCT/AU2012/001321 (TECH ID:2012-009) with royalties paid. The remaining authors have no conflicts of interest.

## Funding acknowledgments

This work was supported by National Health and Medical Research Council (NHMRC) Program Grant (10915693) to SFB, IES and CAR and by an anonymous philanthropic gift for SUDEP research to SFB and IES. CAR would like to acknowledge the CURE foundation for support. CS and IES are recipients of a National Health and Medical Research Council Practitioner Fellowships (#1154992 to CS and #1104831 to IES) and Senior Investigator Fellowship to IES (# 1172897). MB was supported by an NHMRC Senior Research Fellowship (#1102971). LEB acknowledges the support of an Australian Government Research Training Program Scholarship. This work was also made possible through the Victorian State Government Operational Infrastructure Support and Australian Government National Health and Medical Research Council (NHMRC) independent research Institute Infrastructure Support Scheme (IRIISS).

